# Structure and dynamics of human disease-complication network

**DOI:** 10.1101/2021.12.13.472342

**Authors:** Xiong-Fei Jiang, Long Xiong, Ling Bai, Jie Lin, Jing-Feng Zhang, Kun Yan, Jia-Zhen Zhu, Bo Zheng, Jian-Jun Zheng

## Abstract

A complication is an unanticipated disease arisen following, induced by a disease, a treatment or a procedure. We compile the Human Disease-Complication Network from the medical data and investigate the characteristics of the network. It is observed that the modules of the network are dominated by the classes of diseases. The relations between modules are unveiled in detail. Three nontrivial motifs are identified from the network. We further simulate the dynamics of motifs with the Boolean dynamic model. Each motif represents a specific dynamic behavior, which is potentially functional in the disease system, such as generating temporal progressions and governing the responses to fluctuating external stimuli.

**Author summary:** Advances in molecular biology lead to a new discipline of network medicine, investigating human diseases in a networked structure perspective. Recently, clinical records have been introduced to the research of complex networks of diseases. An important available medical dataset that has been overlooked so far is the complications of diseases, which are vital for human beings. We compile the Human Disease-Complication Network, representing the causality between the upstream diseases and their downstream complications. This work not only helps us to comprehend why certain groups of diseases appear collectively, but also provides a new paradigm to investigate the dynamics of disease progression. For clinical applications, the investigation of complications may yield new approaches to disease prevention, diagnosis and treatment.

## Introduction

Advances in molecular biology lead to a new discipline of network medicine, which has multiple potential biological and clinical applications to human disease [1–5]. Much attention has been paid to molecular networks, including protein interaction networks [6–9], regulatory networks [10], metabolic networks [11–14], RNA networks [15–17], and kinase-substrate networks [18], whose nodes are molecules linked to each other by physical interactions. In parallel, an increasing number of studies focus on phenotypic networks, including co-expression networks [19, 20], and genetic networks [21–23].

The highly interconnected property of the interactome suggests that diseases are not independent of one another, whether at the molecular level or phenotypic level. From psychiatric patient records, comorbidities are determined by the co-occurring frequency of disease pairs in patients more often than expected [24]. Khan et. al. investigate the chronic disease progression pattern of Type 2 Diabetes by the comorbidity network drawn from the Australian healthcare records [25]. The phenotypic disease network with comorbidity patterns is extracted from more than 30 million medicare records, and particularly the dynamic progression patterns are investigated [26]. Further, the genetic overlap between diseases is estimated with the comorbidity correlation, which is extracted from the disease history of 1.5 million patients [27]. Similarly, Zhou et. al. observe that the symptom-based similarity between two diseases positively correlates with the number of shared genes from the human disease network [28]. The projection of such network-based dependencies between disease genes and disease phenotypic features has brought the concept of diseasome [1, 29–31].

To go beyond the global features, the motif is utilized to characterize the local properties of those biological networks. The pioneering work by Milo et al. identifies motifs in networks from biochemistry, neurobiology, ecology, and engineering [32]. Then motifs have been observed in many biological networks [33–35], and their dynamics have been investigated further [34, 36].

A critical medical dataset that has been overlooked so far is the complications of diseases, which are vital for patients in clinical practice. For example, complictions may affect clinical decisions for physicians in certain in circumstances [37]. A complication is an unanticipated disease arisen following, induced by a disease, a treatment or a procedure. Diseases form a networked structure with their complications. For example, COVID-19 generates numerous complications, such as venous thromboembolism and acute kidney injury, etc [38]. We compile the Human Disease-Complication Network, representing the causality between the upstream diseases and their downstream complications. We systematically investigate the structure of the network, and pay attention to the disease modules. Taking into account the complexity of network dynamics, we further study the motifs, with which the dynamics can be depicted in the disease system. This work not only helps us understand how different medical subdisciplines organize, but also provides a comprehensive understanding of why certain groups of diseases appear collectively.

## Materials and methods

### Materials

We collected data from the Clinical Medicine Knowledge Database, including 6715 diseases. The complications of diseases are extracted from the descriptions of diseases in the database. A node denotes a disease, and a directed link from the *i*-th node to the *j*-th node is drawn if the *i*-th disease generates the *j*-th complication. Then, we compile the directed Human Disease-Complication Network (HDCN), containing 3471 nodes and 9164 directed links. The HDCN is represented by the adjacency matrix 𝔾 = [*G*_*ij*_], where *G*_*ij*_ = 1 if there is a directed link pointing from the *i*-th node to the *j*-th, otherwise *G*_*ij*_ = 0. The giant connected component includes 3390 nodes and 9116 directed links. The data is available in the supporting information.

## Methods

### *k***-core**

The *k*-core analysis is able to uncover a nucleus set of nodes, i.e., a set of nodes with a high degree connected to each other. It has been widely used in networks to identify the kernels more robustly than simply through the ranking of centrality measures [39, 40]. The *k*-core of a network consists of nodes *i* with the degree *k*_*i*_ *k*, and the *k*-core could be extracted by the iterative removal of all nodes *i* with degrees *k*_*i*_ *< k* when *k >* 0.

### Motif

The motifs are defined as local patterns occurring in the real network significantly more frequently than in randomized networks with the same degree sequence [32, 34]. For any given network, the occurrence number *N*_*g*_ of the *g*-th connected subsets is related to the network size and the degree distribution. For measuring the statistical significance, the randomized ensemble of networks is generated as a null model [32]. The statistical significance *Z* score is defined as

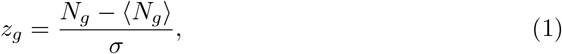

where *N*_*g*_ is the appearance number of the *g*-th subset in the real HDCN, ⟨*N*_*g*_⟩ and *σ* are the average appearance number and standard deviation in the randomized ensemble of networks. The motifs are those subsets with significantly higher frequency in the real network than randomized ones, measured by the *Z* score. In this paper, we mainly focus on the 3-node and 4-node connected subsets.

### Boolean dynamics

The Boolean dynamics is a common model in gene regulatory networks and signal transduction networks [41, 42]. Each node in a Boolean network represents a sub-cellular component such as protein, gene, transcription factor or metabolite. The states of input nodes *i* are described by a binary value *X*_*i*_. *X*_*i*_ = 1 represents that the component *i* is active or expressed, *X*_*i*_ = 0 means that it is inactive or not expressed. The state of node *j* at time *t* + 1, *X*_*j*_(*t* + 1), is determined by a logic operation together with the current state of its upstream regulators *X*_*i*_(*t*). The logic operation is the Boolean update function, denoted by the logic operators or a weighted sum of the inputs to an activation threshold. The output nodal dynamics is described by a differential equation like

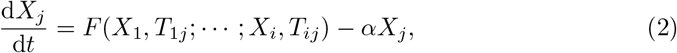

where *F* (*X*_1_, *T*_1*j*_; ; *X*_*i*_, *T*_*ij*_) is the Boolean update function.

Here, we introduce the Boolean dynamic model to simulate the complication progression in the HDCN. The disease progression in the HDCN is compared to other biological processes, such as gene regulatory networks. A complication is generated by the collective effect of upstream diseases, and it will presumably self-cure after the upstream diseases are cured. It is reasonable to utilize the Boolean dynamic model to investigate the complication dynamics. Therefore, *X*_*i*_ represents the activation of the *i*-th upstream disease, *T*_*ij*_ is the activation threshold of the *i*-th disease to the *j*-th complication, *α* is the lifetime of the cured disease, and *F* (*X*_1_, *T*_1*j*_; ; *X*_*i*_, *T*_*ij*_) denotes the summarized effect of overall *i* upstream diseases on the *j*-th complication.

## Results

Qualitatively, the HDCN is formed by very few disconnected components and a large giant connected component (see Fig 1), including approximately 93.64% of all diseases. The majority of the nodes exhibit low clustering coefficients (0 *< c*_*i*_ *≤* 0.1), representing a sparse local structure, while the minority demonstrate high clustering coefficients (0.1 ≤ *c*_*i*_ *<* 1.0), indicating a dense local structure. The disease-complication relations in the HDCN are specific, 16%, 32% and 21% of the diseases generate 1, 2 and 3 complications respectively, while only 31% diseases cause more than 3 complications. Vice versa, 73% of the diseases are only caused by a single disease, 10% are caused by two. We calculate the probability distribution of the degrees, which are distinguished between in-degree, i.e. the number of edges pointed to some vertex, and out-degree, i.e. the number of edges pointing away from it. As shown in Fig 2, the probability distribution of out-degree exhibits an approximative power-law, while the one of in-degree decays in an exponentially. The out-degree distribution of the HDCN indicates that most diseases generate only a few other diseases, whereas a few phenotypes such as Scleroderma (*k*^*out*^ = 23) and Renal Damage due to Hyperthyroidism (*k*^*out*^ = 19) generate a large number of distinct diseases. The in-degree distribution decays faster than the out-degree distribution, but still significantly deviates from the Poisson distribution expected for a random graph.

**Fig 1.**
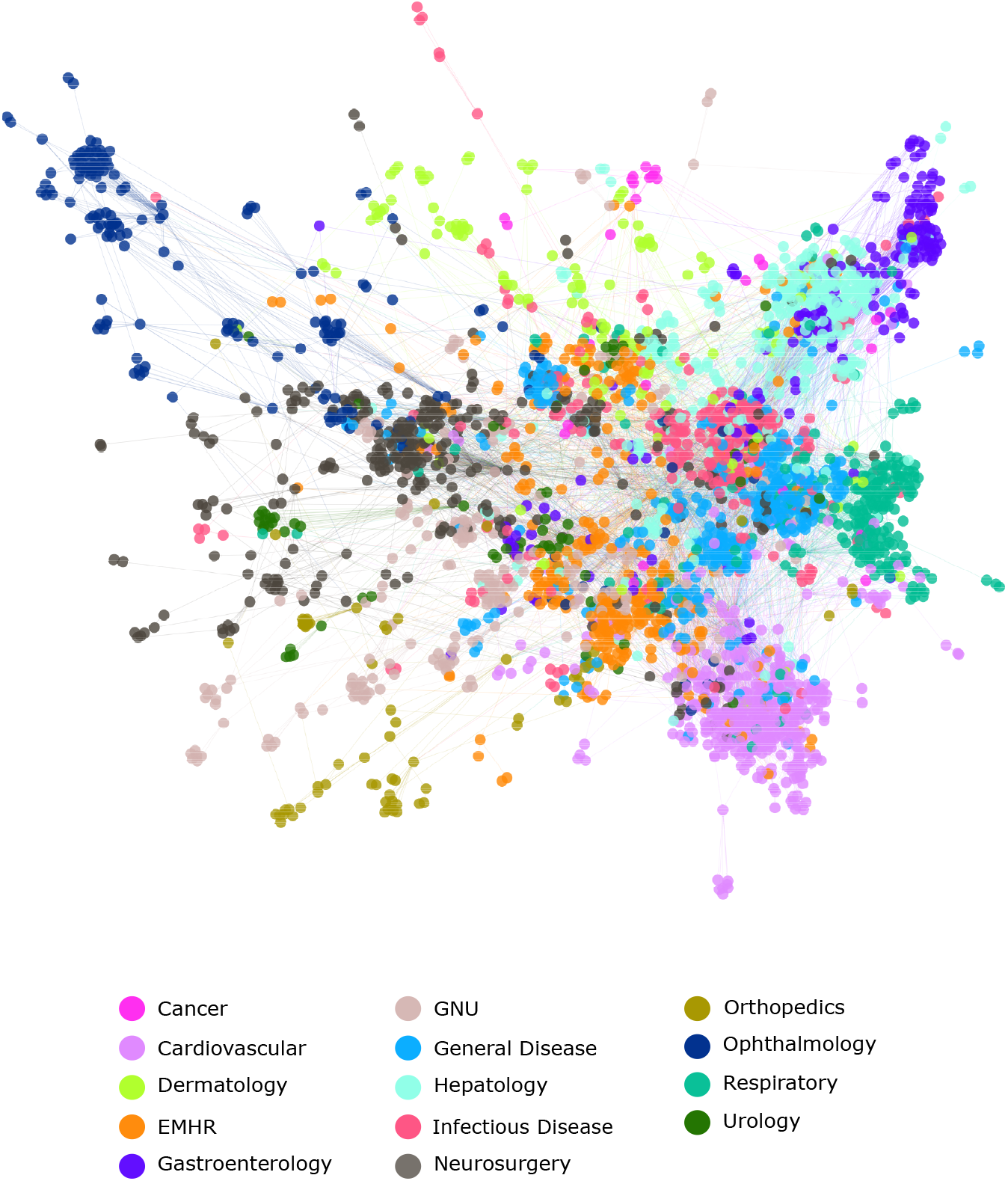
Human Disease-Complication Network. Each node corresponds to a distinct disease, colored based on the disease class to which it belongs. Names of the 14 disease classes are indicated on the bottom.

**Fig 2.**
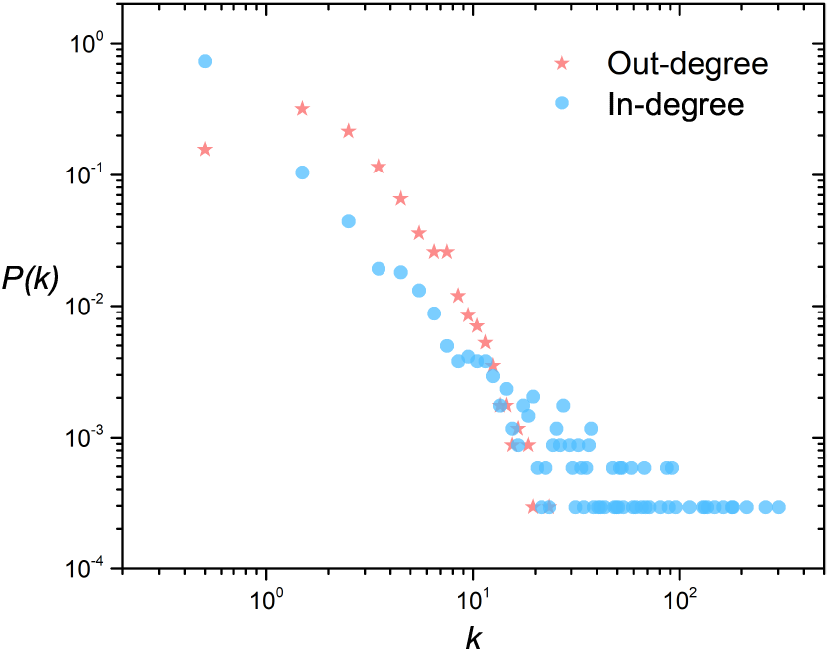
Degree distribution. The out-degree and in-degree distributions *P* (*k*) of the HDCN. In-degree is the number of edges pointed to some vertex, and out-degree is the number of edges pointing away from it. the probability distribution of out-degree exhibits an approximative power-law, while the one of in-degree decays in an exponentially.

In the HDCN, a disease presents a streaming structure: generating complications, and as a complication caused by other diseases. Therefore, we introduce in-degree *k*^*in*^ and out-degree *k*^*out*^ to quantitatively categorize the diseases into these three structural levels, i.e., upstream, intermediate and downstream. The diseases with highest *k*^*out*^ and *k*^*in*^ are listed in Table 1. A node with high *k*^*out*^ and low *k*^*in*^ is an upstream disease such as Acute Lymphoblastic Leukemia (*k*^*out*^ = 16, *k*^*in*^ = 1), since it could lead to other diseases yet be hardly produced by others. On the contrary, a node with low *k*^*out*^ and high *k*^*in*^ is a downstream disease such as Pneumonia (*k*^*out*^ = 0, *k*^*in*^ = 181), which almost results from the other diseases, nevertheless could hardly produce others. A node, which could generate other diseases and be produced by others, is an intermediate disease. In this case, the terms of upstream, intermediate and downstream are not absolute, but relative. Although it is less likely to do so, a downstream disease could cause complications. Qualitatively, diseases with the highest *k*^*in*^ are usually general diseases, while nodes with the highest *k*^*out*^ are frequently associated with specific organs.

**Table 1.** Diseases with the highest out-degree *k*^*out*^ and in-degree *k*^*in*^ in the HDCN. RDH: Renal Damage due to Hyperthyroidism, PPNS: Pediatric Primary Nephrotic Syndrome, ALL: Acute Lymphoblastic Leukemia, PNH: Paroxysmal Nocturnal Hemoglobinuria, SRHD: Senile Rheumatic Heart Disease, CACCR: Cardiac Arrest and Cardiopulmonary-Cerebral Resuscitation, CAHD: Coronary Atherosclerotic Heart Disease, SSH: Spontaneous Subarachnoid Hemorrhage, SIE: Subacute Infective Endocarditis, ATH: Alimentary Tract Hemorrhage.

| Disease | $k^{out}$ | Disease | $k^{in}$ |
| --- | --- | --- | --- |
| Scleroderma | 23 | Shock | 302 |
| RDH | 19 | Heart Failure | 261 |
| Chronic Cor Pulmonale | 18 | Arrhythmia | 211 |
| Hyperthyroidism | 18 | Pneumonia | 181 |
| PPNS | 18 | Septicemia | 179 |
| Viral Hemorrhagic Fever | 16 | Hypertension | 163 |
| Epidemic Hemorrhagic Fever | 16 | Dehydration | 136 |
| ALL | 16 | Jaundice | 132 |
| PNH | 15 | Ileus | 129 |
| SRHD | 15 | Diarrhea | 111 |
| CACCR | 14 | Myocarditis | 95 |
| Acute Infective Endocarditis | 13 | Diabetes Mellitus | 91 |
| Cholelithiasis | 13 | Ascites | 91 |
| Pediatric Acute Pancreatitis | 13 | Pulmonary Edema | 88 |
| Senile Hyperthyroidism | 13 | Hepatic Cirrhosis | 86 |
| Hepatic Cirrhosis | 12 | ATH | 86 |
| CAHD | 12 | Thrombosis | 68 |
| SSH | 12 | Congestive Heart Failure | 65 |
| Mitral Stenosis | 12 | Pleurisy | 58 |
| SIE | 12 | Myocardial Infarction | 58 |

To quantify the correlation between the in-degree and out-degree of nodes, we compute the Spearman rank correlation coefficient *SC* between these two ranking [43]. A negative value of *SC* = − 0.26 indicates that the in-degree and out-degree in the same node are significantly asymmetric, i.e., the disease connecting with more upstream diseases usually results in fewer downstream complications, and vice versa.

With the *k*-core method [39, 40], we identify a small well-connected nucleus, consisting of 98 diseases with the maximal coreness= 7 (See S1 Table). The components of the nucleus mainly belong to three classes of diseases, i.e., Cardiovascular, Hepatology, and General Diseases. It suggests that these diseases play a crucial role in the progression of complications.

### Disease Modules

Although the HDCN layout is generated without any priori knowledge on disease classes, it is naturally and visibly clustered according to major disease classes. In order to quantitatively understand this clustering nature, we identify the community structure of the HDCN. In a complex network, the community structure is the grouping of nodes into clusters with a high density of internal links, while including a relatively low density of links between clusters [44, 45].

As shown in Fig 1, the diseases in the HDCN form several functional modules dominated by the organs in which the diseases occur. We identify 14 modules, including Orthopedics, Gastroenterology, Cancer, Neurosurgery, Dermatology, Urology, Cardiovascular, Respiratory, Ophthalmology, Infectious Disease, General Disease, Hepatology, Gynecology-Nephrology-Urology (GNU) and Endocrine-Metabolic-Hematology-Rheumatology (EMHR). Hepatology includes Hepatobiliary, Pancreatic and Splenic Diseases. It should be noted that the Cancer module only includes a small fraction of cancers. The majority of cancers are distributed in the related organs, rather than form a single module as in the gene-disease network [30]. This difference is rooted in that the cancers in the former one usually cause complications in the related organs, while the latter one may share the same genes and generate dense connections between each other. Gynecology, Nephrology and Urology are grouped into a single module, since all three belong to the genitourinary system in which the related organs are physiologically close and interlinked by the blood supply and some meatus. In a similar vein, the diseases in Endocrine, Metabolic, Hematology and Rheumatology form a single module, which has a global effect on the whole human disease system.

To quantify the influence of modules in the progression of complications, we define the complicating ability of the *p*-th module as

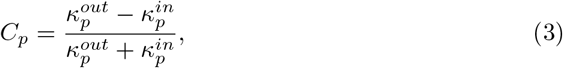

where 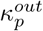 and 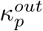 are the total out-links and in-links of the *p*-th module connecting to other modules. The high value of *C*_*p*_ suggests that the module can trigger downstream complications with high possibility, while the high absolute value of negative *C*_*p*_ means that the module is prone to be caused by upstream diseases. As shown in Table 2, Orthopedics, Dermatology, and Cancer are the modules with the highest *C*_*p*_. Therefore, these three are the most influential disease modules, which can generate the downstream diseases with great capability. Meanwhile, General Disease, Cardiovascular and Infectious Disease are the modules with the lowest *C*_*p*_. It implies that they can be triggered by other diseases easily.

**Table 2.** Modules in the HDCN. Here *N* is the number of diseases in modules, *m* is the number of links connecting nodes within modules, *M*^*out*^ and *M*^*in*^ are the total numbers of out-links and in-links connecting nodes in other modules. Hepatology includes Hepatobiliary, Pancreatic and Splenic Diseases. GNU: Gynecology-Nephrology-Urology, EMHR: Endocrine-Metabolic-Hematology-Rheumatology.

| | $N$ | $m$ | $M^{out}$ | $M^{in}$ | $r$ |
| --- | --- | --- | --- | --- | --- |
| Orthopedics | 77 | 84 | 29 | 11 | 0.45 |
| Gastroenterology | 184 | 279 | 177 | 179 | -0.01 |
| Cancer | 30 | 37 | 26 | 11 | 0.41 |
| Neurosurgery | 322 | 477 | 231 | 203 | 0.06 |
| Dermatology | 140 | 168 | 145 | 55 | 0.45 |
| Urology | 65 | 72 | 47 | 39 | 0.09 |
| Cardiovascular | 413 | 921 | 280 | 438 | -0.22 |
| Respiratory | 248 | 432 | 255 | 181 | 0.17 |
| GNU | 230 | 341 | 166 | 149 | 0.05 |
| Hepatology | 353 | 670 | 359 | 282 | 0.12 |
| Ophthalmology | 177 | 267 | 61 | 50 | 0.10 |
| EMHR | 306 | 469 | 369 | 260 | 0.17 |
| Infectious Disease | 297 | 445 | 254 | 342 | -0.15 |
| General Disease | 356 | 531 | 274 | 473 | -0.27 |

To further unveil the causal relations between the disease modules, we define the density of directed links from the *i*-th to the *j*-th module, 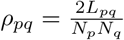, where *L*_*pq*_ is the number of directed links from the *p*-th module to the *q*-th module, *N*_*p*_ is the number of diseases in the *p*-th module, and *ρ*_*pq*_ *≠ ρ*_*qp*_. The average density of links for the entire network is 6.8 × 10^−4^, and the one within the modules is 2.0 × 10^−2^. We set the average density of links between the modules, 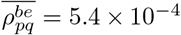, as the threshold to determine the backbone of the HDCN. In other words, only the links whose *ρ* is higher than 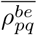 remain.

Fig 3 outlines the relations and directions between the disease modules. EMHR and Dermatology are the two modules with the strongest influence. For example, EMHR generates almost half of the complication modules, including Cancer, Hepatology, GNU, Cardiovascular, and General Disease. On the other hand, Infectious Disease and General Disease modules are the two most attractive modules. Infectious Disease is possibly caused by Dermatology, Respiratory, Gastroenterology, Hepatology, and Neurosurgery with high possibilities. Ophthalmology and Orthopedics modules are relatively independent in the HDCN. Ophthalmology only causes Neurosurgery, and Orthopedics only leads to General Diseases. It implies that Ophthalmology and Orthopedics have a specific impact on the complication progression. Gastroenterology and Hepatology have the strongest bi-directed causalities in the HDCN since both of them belong to the digest system. Besides, EMHR may lead to Cardiovascular, and Dermatology may result in Infectious Disease with high probabilities, respectively. Notably, Cancer module does not have a very high complication effect on other modules. The ultimate reason is that there are only 32 cancers included in Cancer module (40 diseases in total), and most cancers are classified into the corresponding modules of organs.

**Fig 3.**
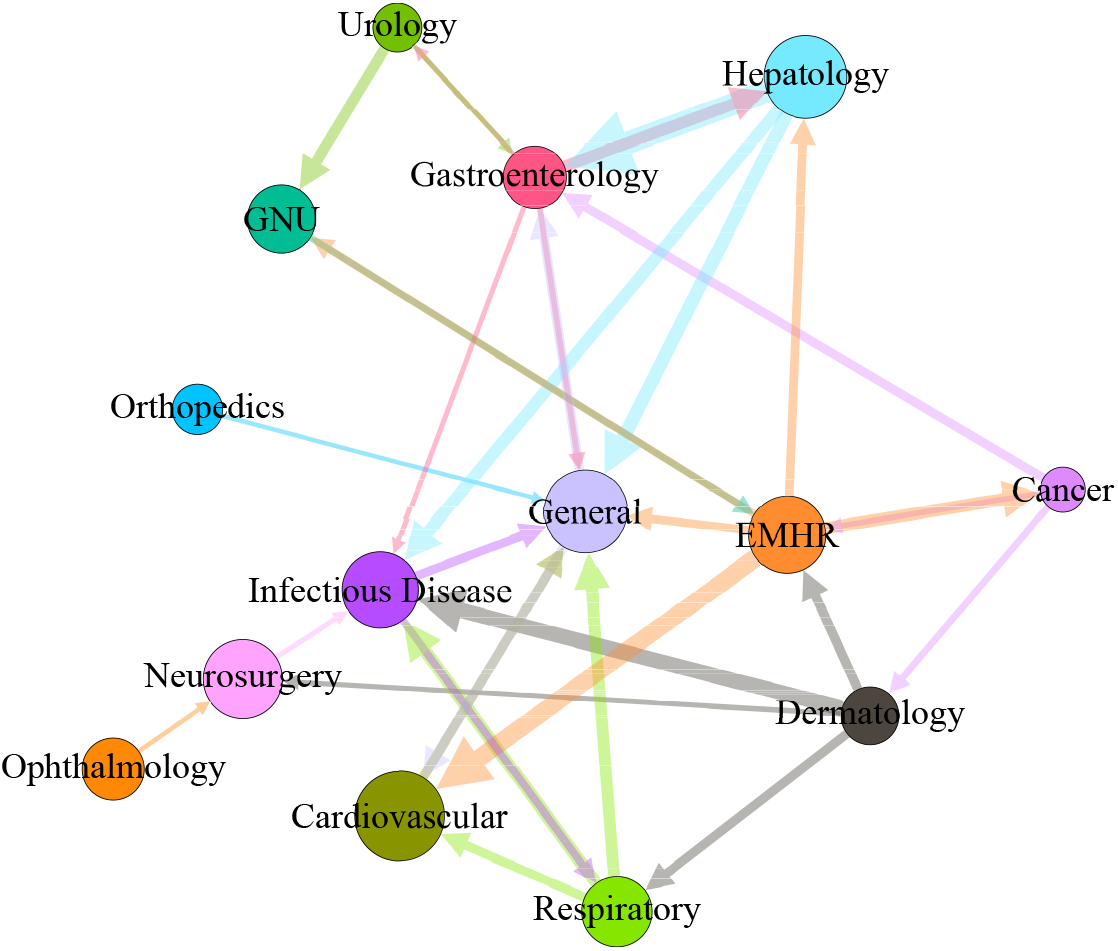
Causal connections between disease modules. The node size represents the number of diseases in the module, while the width of the line suggests the density of the connection between two modules.

### Motif

To mitigate the complexity of the HDCN, we investigate the local interactions among small connected subsets, i.e., motifs. The statistical significance of the network motifs is evaluated by comparison with randomized networks having the same characteristics as the HDCN. We identify three kinds of motifs, i.e., the feedforward loop, the bi-fan, and the overlapping feedforward loop, as shown in Table 3.

**Table 3.**
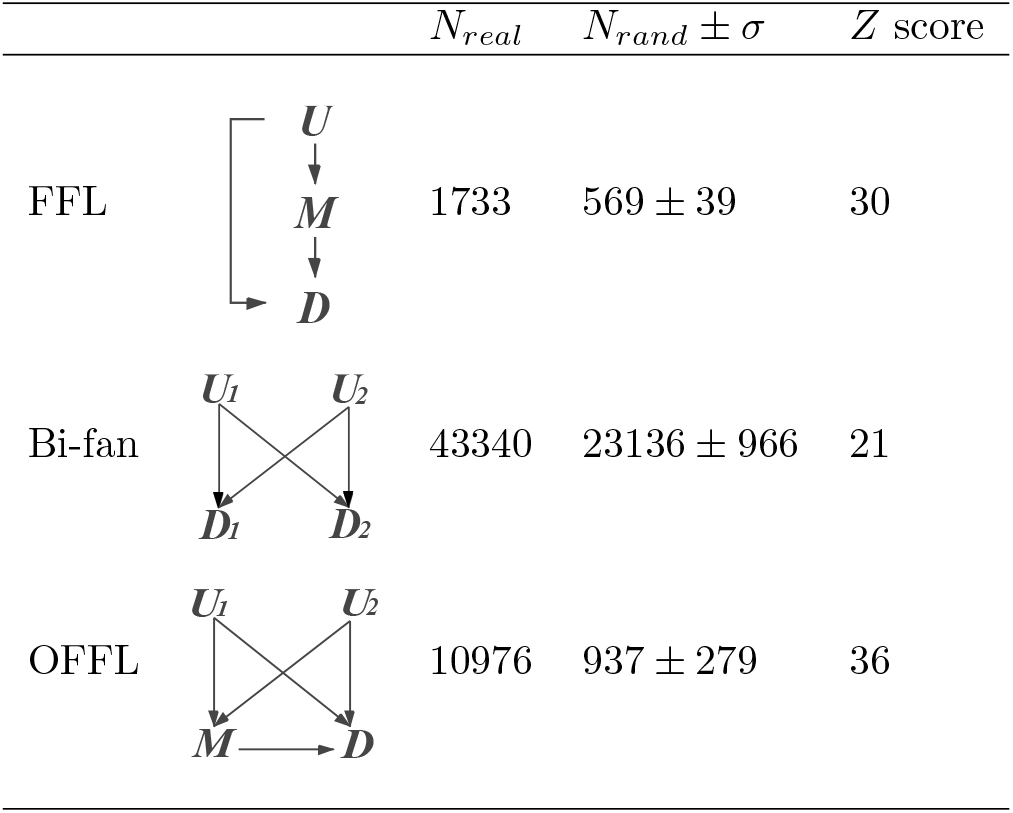
Network motifs detected in the HDCN. For each motif, the numbers of appearances in the real network *N*_*real*_, and in the randomized networks *N*_*rand*_ ± *σ* are listed. The *P* value of all motifs is *P <* 0.01, as determined by comparsion to 1000 randomized networks. The *Z* score= (*N*_*real*_ − *N*_*rand*_) */σ*, as a measure of statistical significance. *U* : upstream disease, *M* : mediated disease, *D*: downstream disease.

| | | $N_{real}$ | $N_{rand} \pm \sigma$ | $Z$ score |
| --- | --- | --- | --- | --- |
| FFL | | 1733 | $569 \pm 39$ | 30 |
| Bi-fan | | 43340 | $23136 \pm 966$ | 21 |
| OFFL | | 10976 | $937 \pm 279$ | 36 |

### Feedforward loop

The feedforward loop (FFL) is formed by three diseases: *U* (upstream disease) generates *M* (mediated disease) and *D* (downstream disease), while *M* also causes *D*, as illustrated in Table 3. For instance, pediatric food allergy (*U*), allergic purpura (*M*) and arrhythmia (*D*) consist of a typical FFL. To deeply understand the medical function of the FFL, we search all compositions of diseases in the motif. The FFL appears a total 1733 times in the HDCN. Thereinto, *U, M* and *D* nodes contain 722, 274, 193 unique diseases, respectively. The diseases with top 10 appearing times in the three nodes are displayed in S2 Table. Most of the diseases in *U* belong to the upstream diseases, the diseases in *M* are generally the intermediate diseases, while the diseases in *D* tend to be the downstream diseases.

We calculate the average in-degree 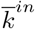 and out-degree 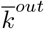 for *U, M, D* nodes in the FFL, listed in Table 4. It is observed that 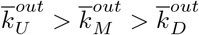, whereas 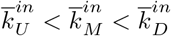. These results quantitatively confirm that *U, M* and *D* nodes in the FFL tend to be the upstream, intermediate and downstream diseases, respectively.

**Table 4.** Average in-degree *k*^*in*^ and out-degree *k*^*out*^ of nodes in the motifs. The values are computed from the weighted average. *U* : upstream disease, *M* : mediated disease, *D*: downstream disease.

| FFL | $U$ | $M$ | $D$ | |
| --- | --- | --- | --- | --- |
| $\bar{k}^{out}$ | 8.55 | 6.59 | 1.41 | |
| $\bar{k}^{in}$ | 1.90 | 52.86 | 124.64 | |
| Bi-fan | $U_1$ | $U_2$ | $D_1$ | $D_2$ |
| $\bar{k}^{out}$ | 9.18 | 8.89 | 1.12 | 0.90 |
| $\bar{k}^{in}$ | 3.12 | 1.77 | 16.94 | 156.90 |
| OFFL | $U_1$ | $U_2$ | $M$ | $D$ |
| $\bar{k}^{out}$ | 7.82 | 7.58 | 5.04 | 0.51 |
| $\bar{k}^{in}$ | 1.75 | 1.84 | 148.25 | 204.59 |

### Bi-fan

The bi-fan is comprised of four nodes, classified into two layers between which there exist overlapping interactions, as demonstrated in Table 3. For instance, group A streptococci (*U*_1_), brucellosis (*U*_2_), pneumonia (*D*_1_) and endocarditis (*D*_2_) consist of a bi-fan. We search all diseases appeared in the bi-fan as well. *U*_1_, *U*_2_, *D*_1_, *D*_2_ nodes encompass 1399, 1401, 317 and 327 unique diseases, respectively. The diseases with top 10 appearing times in the four nodes are listed in S3 Table. Most of the diseases in *U*_1_ and *U*_2_ belong to upstream diseases, while *D*_1_ and *D*_2_ tend to be downstream diseases. Hence, we call *U*_1_ and *U*_2_ as upstream disease layer, while *D*_1_ and *D*_2_ as downstream disease layer. 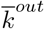 and 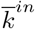 of four nodes in the bi-fan are shown in Table 4.

The average in(out)-degree of *U*_1_ and *U*_2_ are almost the same, while the average in(out)-degrees of *D*_1_ and *D*_2_ are similar, since the structure of the bi-fan is symmetric. Then 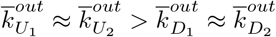, while 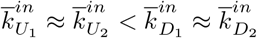. The results ascertain that *U*_1_ and *U*_2_ nodes tend to be the upstream diseases, meanwhile *D*_1_ and *D*_2_ nodes would be the downstream diseases in the bi-fan.

### Overlapping feedforward loop

Further, we identify a novel motif called overlapping feedforward loop (OFFL), which has not been observed in other networks. As shown in Table 3, the OFFL consists of two FFLs, sharing *M* and *D* nodes. For instance, viral hemorrhagic fever (*U*_1_), cholelithiasis (*U*_2_), septicemia (*M*) and jaundice (*D*) form an OFFL. There are 548, 546, 132 and 97 unique diseases included in *U*_1_, *U*_2_, *M* and *D* nodes respectively. The diseases with top 10 appearing times in the four nodes are listed in S4 Table. Most of the diseases in *U*_1_ and *U*_2_ nodes can be categorized as upstream, whilst diseases in *M* as intermediate, and diseases in *D* as downstream.

Table 4 displays 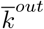 and 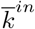 for the four nodes in the OFFL. The average in(out)-degrees of *U*_1_ node are almost the same as those of *U*_2_ node. *U*_1,2_ nodes have a high 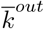 with low 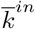, while the *D* nodes have a high 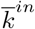 with low 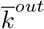. Besides, 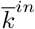 and 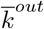 of *M* are intermediate between *U*_1,2_ and *D*. It quantitatively supports that *U*_1_ and *U*_2_ in the OFFL tend to be upstream diseases, *M* are supposed to be intermediate diseases, while *D* would be downstream diseases. This result consolidates the earlier proposition that OFFL is formed with two FFLs, which share the same intermediate and downstream diseases.

### Dynamics

The frequency of motifs appeared significantly higher than in a random network, implying that the motifs may represent medical functions. The dynamics of motifs offers an insight to understand the progression of diseases further. Here we introduce the Boolean dynamic model to simulate the dynamics of the motifs in the HDCN, and only consider the effect of several typical mediated pathways on the dynamics in this paper.

### Feedforward loop

In the FFL motif, the effect of the upstream disease *U* is transferred to the downstream disease *D* via two pathways, a direct one of *U* → *T* and a mediated one of *U* → *M* → *D. U* and *M* may act in an AND-gate or OR-gate manner to control *D*. For the AND-gate, the differential equations are written as

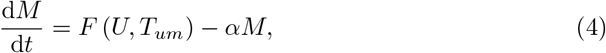

and

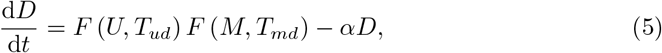

where *F* (*u, T*) = (*u/T*) */* (1 + (*u/T*)). In the remainder of this paper we will adopt this function for all AND-gate, unless stated otherwise. The dynamic behaviors of the AND-gate are shown in Fig 4.a for different parameters. The upstream disease *U* generates the intermediate disease *M*, then *U* and *M* jointly stimulate the downstream disease *D*. When the activation of *U* is transient, the morbidity of *D* is low. In other words, the stimulus can not be transduced through the circuit easily. The downstream disease *D* will be activated only if *U* is a persistent stimulus. Meanwhile, *M* will decrease rapidly, once the stimulus of *U* vanishes. The FFL motif may resist stochastic perturbations in the environment, and prevent the wild disease progression in the disease system. In fact, the FFL is also a common motif detected in transcriptional gene regulation networks and neuronal connectivity networks [34]. Therefore, the AND-gate of the FFL is probably a general mechanism to protect biological functions.

**Fig 4.**
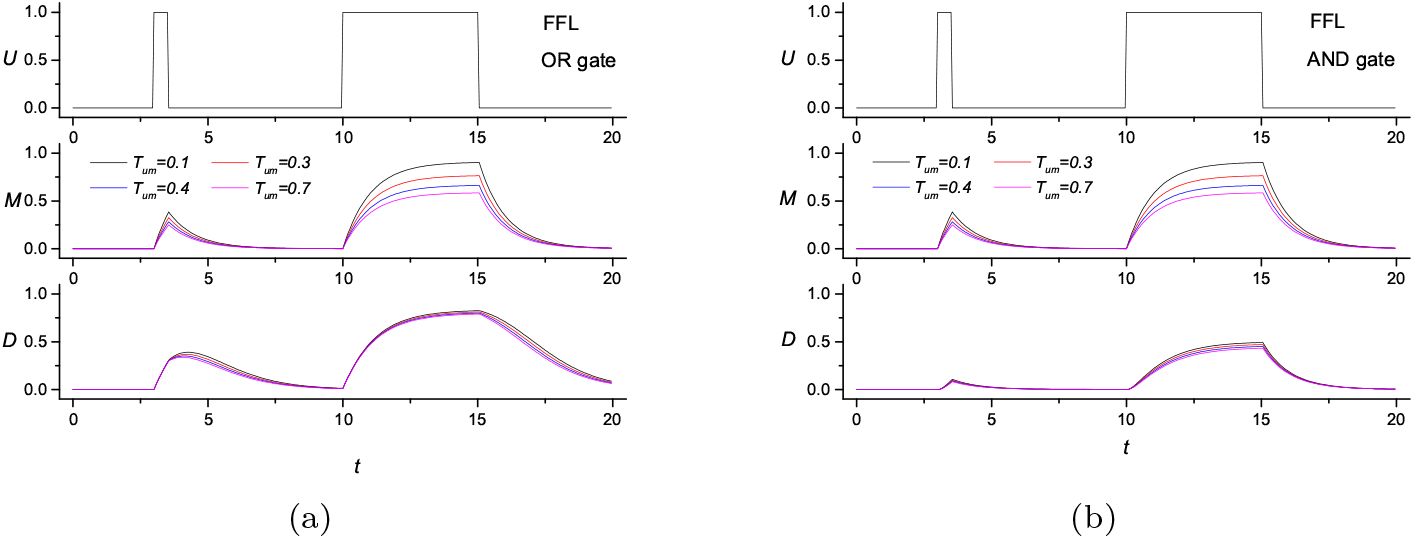
Dynamic features of the feedforward loop motif. *T*_*ud*_ = 0.5, *T*_*wd*_ = 0.3, and *α* = 1.0.

For the OR-gate, Eq 5 is substituted by 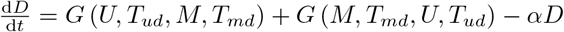, where

*G* (*u, T*_*u*_, *v, T*_*v*_) = (*u/T*) */* (1 + (*u/T*_*u*_) + (*v/T*_*v*_)). If not stated otherwise, the function will be adopted for all OR-gate. The results are exhibited in Fig 4.b. In the OR-gate, *U* and *M* can stimulate the downstream disease *D* independently. The response rate of *D* in the OR-gate is higher than the one in the AND-gate. In contrast, the relaxation rate of *D* in the OR-gate is lower than that in the AND-gate. *D* would be generated even if the stimulus of *U* is transient, since the causing pathway of *U → D* and *M →D* are independent. Besides, the relaxation time of *D* in the OR-gate is much longer than that in the AND-gate when the stimulus of *U* vanishes. The pathway of *M →D* would enhance the morbidity of *D*, whenever the stimulus of *U* is persistent or transient. Hence the circuit of OR-gate may aggravate the disease progression.

### Bi-fan

In the bi-fan motif, upstream diseases *U*_1,2_ have two affecting pathways, *U*_1_ → *D*_1,2_ and *U*_2_ → *D*_1,2_. Likewise, *U*_1_ and *U*_2_ may act in an AND-gate or OR-gate manner to control *D*_1,2_.

For the OR-gate, both *D*_1,2_ could be activated by *U*_1_ and *U*_2_ independently, i.e., *U*_1_ → *D*_1,2_ and *U*_2_ → *D*_1,2_. The differential equations of the bi-fan are written as

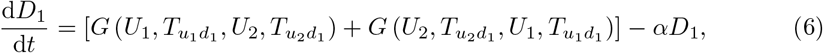

and

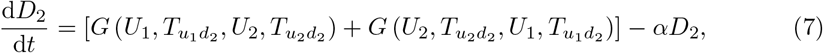

As shown in Fig 5, for the transient signals, if there is only one upstream disease (*U*_1_ or *U*_2_) with a transient signal, the downstream diseases can not be activated. Only when *U*_1_ and *U*_2_ co-occur simultaneously will the morbidity of *D*_1_ and *D*_2_ be high. To some extent, the FFL motif provides a temporal mechanism to prevent the wild disease progression of complications, while the bi-fan motif offers a spatial protection mechanism. Intriguingly, the dynamics of *M* with the type-1 stimulus are different from those with the type-2 stimulus. The latter one is sensitive to the variation of 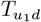, whereas the former is not. Since *U*_1_ remains persistent in type-2 stimulus, while it becomes transient in type-1 stimulus Hence, *D*_1,2_ nodes response sensitively to the persistent stimulus.

**Fig 5.**
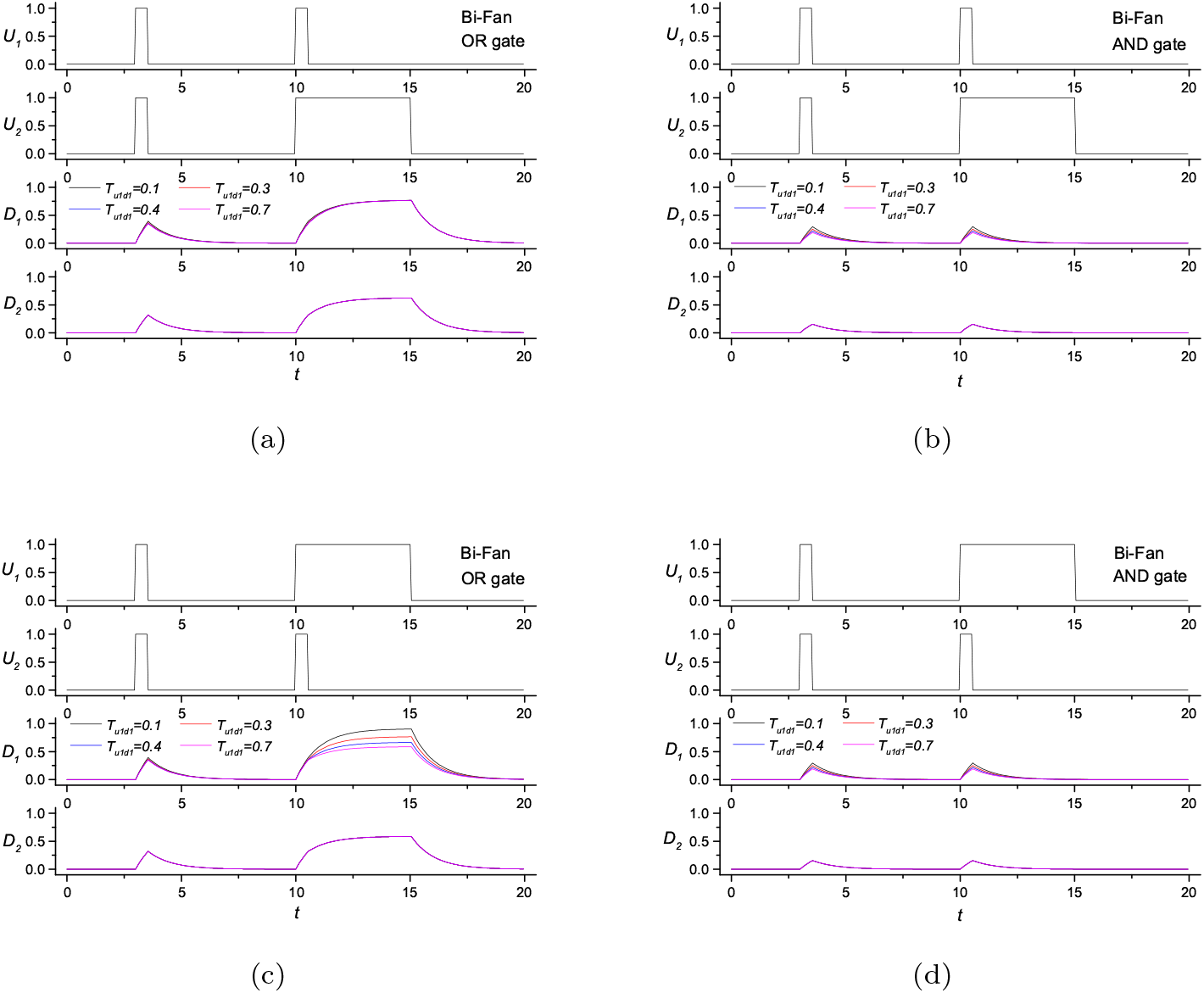
Dynamic features of the bi-fan motif. 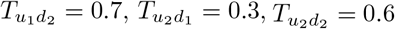, and *α* = 1.0. (a) The OR-gate with type-1 stimulus of *U*_1,2_, (b) The AND-gate with type-1 stimulus of *U*_1,2_, (c) The OR-gate with type-2 stimulus of *U*_1,2_, (d) The AND-gate with type-2 stimulus of *U*_1,2_. Type-1 stimulus: two transient stimuli in *U*_1_, while one transient and one persistent stimuli in *U*_2_. Type-2 stimulus: one transient and one persistent stimuli in *U*_1_, while two transient stimuli in *U*_2_.

For the AND-gate, both *D*_1,2_ should be activated by *U*_1_ and *U*_2_ jointly. Eq 6 and 7 is substituted by 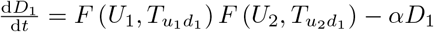 and 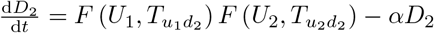, respectively. The results are shown in Fig 5. It is evident that *D*_1_ and *D*_2_ can be activated only if *U*_1_ and *U*_2_ activated simultaneously. The response rates of *D*_1,2_ in the OR-gate are higher than the ones in the AND-gate, while the relaxation rates of *D*_1,2_ in the OR-gate are slower than the ones in the AND-gate.

### Overlapping feedforward loop

The OFFL has a more complicated structure than the other two. For simplicity, we set that *U*_1_ and *U*_2_ always act in an OR-gate manner to control *M* and *D*. However, *M* still acts with *U*_1_ and *U*_2_ in an AND-gate or OR-gate manner to control *D*. We will discuss this AND-gate or OR-gate below.

For a mixed model of AND-gate and OR-gate, the differential equations of the OFFL are written as

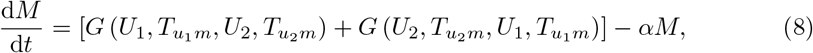

and

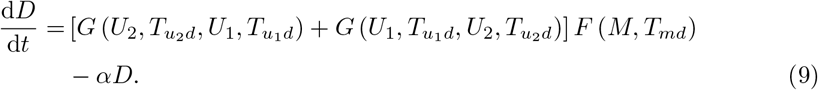

The results are shown in Fig 6. The progression level of *D* disease in the OFFL is significantly lower than the bi-fan, under the same stimulus circumstance of *U*_1_ and *U*_2_. It is counter-intuitive to some extent, since there is an extra pathway from *M* to *D* in the OFFL. Two pathways activating *D*_1,2_ in the bi-fan are direct, and therefore *U*_1_ and *U*_2_ can activate *D*_1,2_ with transient stimulus. By contrast, besides two direct pathways, there is an indirect pathway mediated by *M* in the OFFL. Hence the contribution of pathways of *U*_1_ → *D* and *U*_2_ → *D* are lowered by the pathway of *M* → *D*. Meanwhile, the pathway *M* → *D* should be activated by a persistent stimulus of *U*_1_. Therefore, it is reasonable that the progression level of *D* decreases instead in the OFFL. The loop structure in the OFFL is vital to mediate the disease progression in the HDCN. Similar to the FFL, the AND-gate in the OFFL may reduce the morbidity of complications.

**Fig 6.**
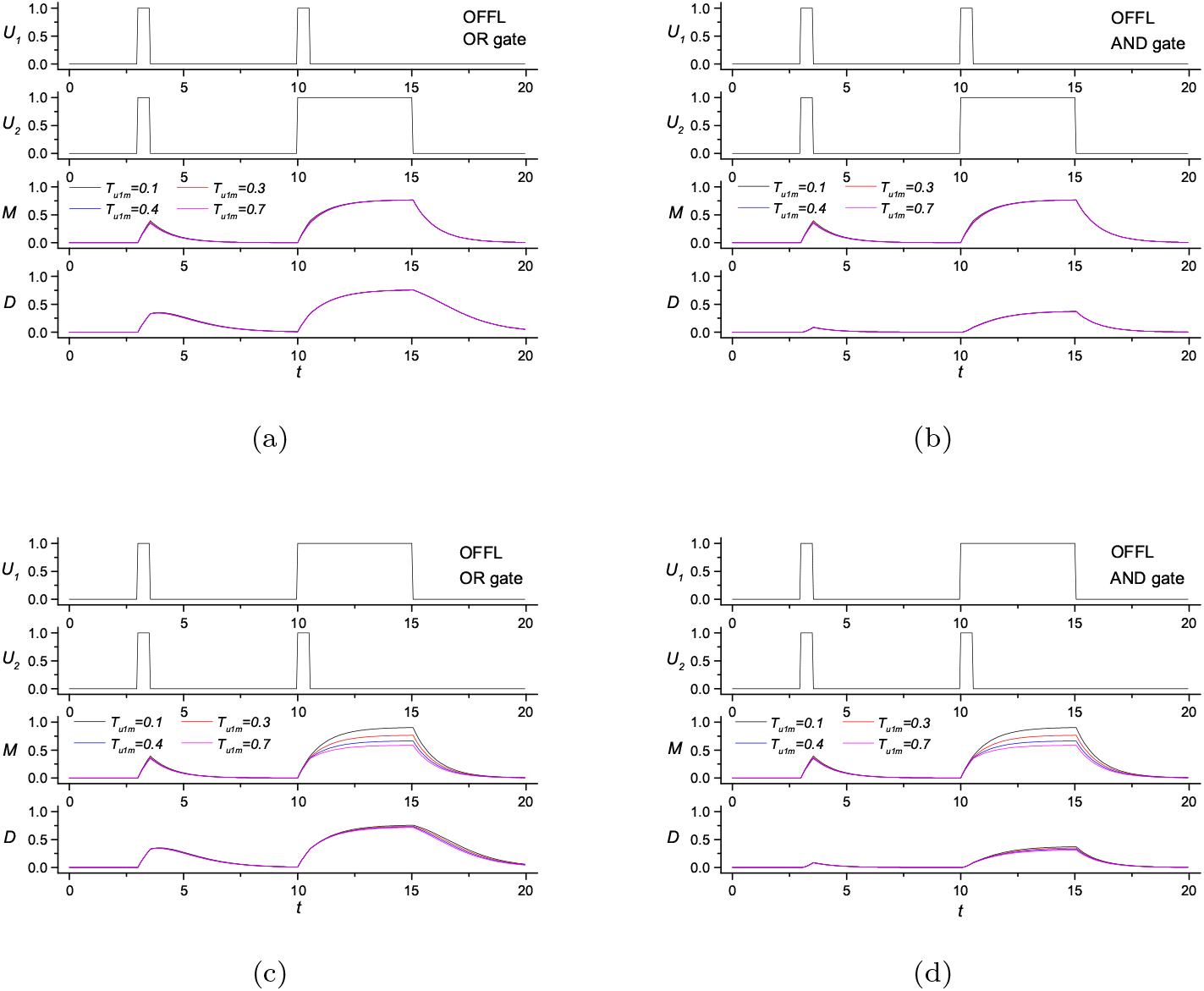
Dynamic features of the OFFL motif. 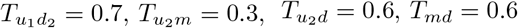 and *α* = 1.0. (a) The OR-gate with type-1 stimulus of *U*_1,2_, (b) The AND-gate with type-1 stimulus of *U*_1,2_, (c) The OR-gate with type-2 stimulus of *U*_1,2_, (d) The AND-gate with type-2 stimulus of *U*_1,2_. Type-1 stimulus: two transient stimuli in *U*_1_, while one transient and one persistent stimuli in *U*_2_. Type-2 stimulus: one transient and one persistent stimuli in *U*_1_, while two transient stimuli in *U*_2_.

For the OR-gate, Eq 9 are written as 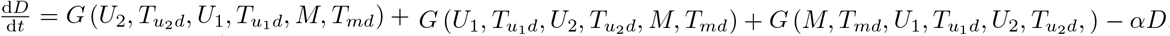. Where *G* (*u, T*_*u*_, *v, T*_*v*_, *w, T*_*w*_) = (*u/T*) */* (1 + (*u/T*_*u*_) + (*v/T*_*v*_) + (*w/T*_*w*_)). The results are shown in Fig 6. Likewise, *D* in the OR-gate presents a dramatic response and longer relaxation time than in the AND-gate. Hence, it is validated that the AND-gate could aggravate the disease progression. Similar to the bi-fan motif, *D* in the OFFL is sensitive to the variation of 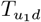.

## Discussion and future work

Complictions may affect clinical decisions for physicians under high pressure and great uncertainty. Singh shows that physicians’ delivery mode decisions (i.e., to perform a vaginal versus a cesarean delivery) are influenced by such a heuristics [37]. If the prior patient had complications in one delivery mode, the physician will be more likely to switch to the other, and likely inappropriate-delivery mode for the subsequent patient, regardless of patient’s indicators. More importantly, this strategy presents small but significantly negative effects on patient health outcomes and increases resource use. There is no clinical reason why the delivery decisions for two separate patients should be causally related to each other. In applications, physicians may overcome the shortcoming of biased decision-making and make appropriate clinical decisions under high pressure, if they have a comprehensive and accurate complication map to evaluate the disease progression quantitatively. Therefore, our work provides a potential framework to aid complex decision-making in clinical practice.

Limitations. There are several limits in the current work. First, we assume nodes are homogeneous, i.e., all diseases have the same capability to cause the downstream complications in Boolean dynamics. However, different diseases have various likelihood of generating complications in practice. For example, COVID 19 has a high likelihood of generating acute kidney injury, a medium likelihood of generating acute liver injury, while a low likelihood of generating immune thrombocytopenia [38]. Second, the update rule of Boolean dynamics in this study is synchronous. However, the complication progression is obviously asynchronous in clinical practice. Third, the dynamical model is only considered in a small scale of motif system.

As part of the future work, we plan to address some of the aforementioned limitations. In particular, we plan to conduct a more comprehensive study on the dynamics of a large network following heterogeneous likelihood of generating complications, with asynchronous update rule. It may initiate a quantitative framework to trace back to the possible original diseases in terms of the current diseases, and predict the progression of complications according to the current diseases. The results from such study will be valuable for diagnosing and developing accurate treatments against complicated diseases, such as COVID-19 and diabetes, which have widespread effects on the whole disease system.

In parallel, many methods have been introduced to the classification of human diseases, such as machine learning [46], integration of phenotypic similarity with genomics [47], pathway-based classification [48] and consensus-based technique [49]. However, contemporary approaches usually do not consider the interactions among diseases [50]. This failure partly comes from the focused nature of medical training, and the reductionist paradigm in modern medicine. To overcome this shortcoming, the network framework is applied to define human disease [29, 51]. In our work, the disease modules clustered in the HDCN may further provide complementary information to classify the human disease more accurately.

## Conclusion

In this paper, the HDCN is constructed from the medical data. We investigate the topological characteristics, including the degree distribution, clustering coefficient and *k*-core of the HDCN. Further, we identify the disease modules which are dominated by the classes of diseases. The relations between modules are unveiled in detail. The 3-node and 4-node motifs are extracted from the HDCN, among which three motifs are significant. We simulate the dynamics of motifs with the Boolean dynamic model. Each motif represents a specific dynamic behavior, which is potentially functional in the disease system, such as generating temporal progressions and governing the responses to fluctuating external stimuli. For clinical applications, the investigation of complications may yield new approaches to disease prevention, diagnosis and treatment.

## Acknowledgments

This work was supported in part by Ningbo Municipal Non-Profit Fund for Applied Research under Grant No. 2019F1033, Ningbo Medical and Health Brand Discipline under Grant No. PPXK2018-05, NNSF of China under Grant No. 11505099, 12175193 and 11775186, Zhejiang Provincial Natural Science Foundation of China under Grant No. LY20G010015.

## Supporting information

**S1 Table. Components of the *k*-core**.

**S2 Table. Diseases with the most occurring times of each node in the FFL motif**.

**S3 Table. Diseases with the most occurring times of each node in the bi-fan motif**.

**S4 Table. Diseases with the most occurring times of each node in the OFFL motif**.

## Data availability

**The data used for this article are available in the supporting information. The source of the directed network is shown in DATA.xlsx, in which the names of nodes are chinese**.

